# Characteristics of spontaneous anterior-posterior oscillation-frequency convergences in the alpha band

**DOI:** 10.1101/2023.09.12.557455

**Authors:** Satoru Suzuki, Marcia Grabowecky, Melisa Menceloglu

## Abstract

Alpha oscillation frequencies vary along the anterior-posterior axis, but they also dynamically converge. We investigated spontaneous anterior-posterior oscillation-frequency convergences while participants rested with their eyes open or closed, by tracking their oscillatory EEG activity in an extended alpha range (5-15 Hz) with appropriate temporal (∽370 ms) and spectral (∽1 Hz) resolutions. Oscillation-frequency convergences were prominent in the alpha band (8-12 Hz) and our analyses revealed three primary characteristics. First, the probability of an additional site joining a frequency convergence increased as more sites frequency-converged, suggesting that synergistic interactions drive anterior-posterior frequency convergences. Second, the oscillatory power at participating sites increased as more sites frequency-converged, suggesting that the synergistic interactions are mediated by regional synchronizations, being boosted by inter-regional frequency matching, facilitating the entrainment of additional regions. Third, frequency convergences generated two opposing phase gradients, posterior-behind and posterior-ahead, potentially mediating directional information flows to and from posterior regions. These gradients became steeper as more sites frequency-converged (while maintaining relatively constant levels of phase consistency), suggesting that frequency convergences increase the directionality of anterior-posterior information flows. Interestingly, when participants closed their eyes, the opposing phase gradients spatially organized, forming posterior-ahead gradients along the midline and posterior-behind gradients within each hemisphere, suggesting that closing eyes streamlines the opposing information flows. Taken together, these results suggest that synergistic interactions drive spontaneous anterior-posterior oscillation-frequency convergences in the alpha band, which may contribute to directional flows of information to and from posterior regions.

## Introduction

Oscillatory neural activity is prevalent in the brain (e.g., Buzsaki, 2006). In particular, inter-regional phase locking of oscillatory activity has been proposed to play a crucial role in controlling neural communications (e.g., Fries, 2005, 2015), spurring much research examining inter-regional phase coherence within and between frequency bands. Most studies measured phase coherence as a period of stable phase difference, Δ*ϕ*(*t*) = *nϕA*(*t*) −*mϕB*(*t*) ∽*constant*, between signals from a pair of brain regions *A* and *B* at frequencies of a ratio, *m*: *n* (where *ϕA*(*t*) and *ϕ****B***(*t*) are instantaneous phases at regions *A* and *B* obtained by Hilbert transforming band-passed signals, convolving with Morlet wavelets, etc.). For example, one would set *m* = *n* = 1 for examining within-frequency phase coherence, *m* = 1 and *n* = 3 for examining cross-frequency phase coherence between *f* and 3 × *f*, and so on. Those studies primarily focused on identifying frequency-specific functional networks organized by within- and/or cross-frequency phase coherence in specific frequency bands. Extensive research using this approach has identified a variety of frequency-specific functional networks and elucidated how they contribute to perceptual, attentional, and cognitive processes (e.g., Lobier et al., 2018; Marzetti et al., 2019; Palva et al., 2005; Palva and Palva, 2012).

Focusing on phase coherence in band-passed signals, however, obscures the fact that oscillation frequencies generally differ across brain regions (e.g., Keitel and Gross, 2016; Mahjoory et al., 2019) as well as they fluctuate over time (e.g., Benwell et al., 2019; Mierau et al., 2017). In order for a pair of regions to stably interact through oscillatory phase locking, their oscillation frequencies need to converge (to the same frequency for within-frequency oscillatory interactions, which was the focus of the current study, or to frequencies of a ratio, *m*: *n*, for cross-frequency oscillatory interactions). Thus, a complementary approach to examining inter-regional oscillatory interactions is to track oscillation frequencies in multiple regions and examine how they dynamically converge. Oscillation frequencies in each region (e.g., an intracranial electrode, EEG/MEG scalp site, inferred source, etc.) can be tracked by applying time-frequency decomposition (e.g., using Morlet wavelets) with appropriate spectral and temporal resolutions, then tracking spectral-power ridges (or stationary-phase points where 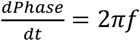 as a function of time (e.g., Amor et al., 2005; Mallat, 2009; Rudrauf et al., 2006). Using this method, previous studies examined spontaneous inter-regional oscillation-frequency convergences in pre-seizure neural activity (e.g., Amor et al., 2005; Rudrauf et al., 2006) as well as frequency convergences to consciously perceived visual flicker during binocular rivalry using frequency-tagged stimuli (Rudrauf et al., 2006).

Our goal here was to uncover macroscopic rules governing spontaneous inter-regional oscillation-frequency convergences during rest using scalp-recorded EEG. Do macroscopic oscillation-frequency convergences occur by coincidence or are they driven? If driven, what mechanisms propel oscillation frequencies in different regions to converge? What functions might inter-regional oscillation-frequency convergences serve? We sought clues to answering these questions by examining the spreading of oscillation-frequency convergences in relation to oscillatory power and phase relations.

We focused on oscillation-frequency convergences along the anterior-posterior axis partly because neural communications between anterior and posterior brain regions have been implicated in a variety of perceptual, attentional, and cognitive processes (e.g., de Pasquale et al., 2012; Gonzalez-Castillo and Bandettini, 2018; Lobier et al., 2018; Marzetti et al., 2019; Mashour et al., 2020; Reinhart and Nguyen, 2019) and partly because consistent anterior-posterior gradients of neuroanatomical and neurophysiological features have been reported including neuron density, myelination, cortical density, functional-connectivity patterns, temporal integration, and oscillation frequency (e.g., Burt et al., 2018; Huntenburg et al., 2018; Mahjoory et al., 2020). We tracked oscillation frequencies in an extended alpha range (5-15 Hz) because they have been implicated in a variety of perceptual, attentional, and memory processes (e.g., Bonnefond and Jensen, 2015; Busch and VanRullen, 2010; Fries, 2015; Harris et al., 2018; Klimesch, 2012; Lobier et al., 2018; Mathewson et al., 2010; Palva and Palva, 2007; Reinhart and Nguyen, 2019), have been shown to influence gamma-band activity (e.g., Bahramisharif et al., 2013; Esghaei et al., 2022), and may organize information flows (e.g., Jensen et al., 2014; Muller et al., 2018; Zhang et al., 2018).

We examined the dynamics of spontaneous anterior-posterior oscillation-frequency convergences in the extended alpha range (5-15 Hz) in representative rest conditions: while participants rested with their eyes closed, rested with their eyes open in a dark room, or viewed a silent nature video. We examined the probabilistic, power, and phase characteristics of inter-regional oscillation-frequency convergences to infer their potential mechanisms and functions. Overall, our results suggest that synergistic interactions drive inter-regional oscillation-frequency convergences in the alpha band, which may facilitate directional flows of information to and from posterior regions.

## Materials and methods

### Participants

Fifty-two Northwestern University students (35 women, 1 non-binary; mean age of 20.8 years, ranging from 18 to 29 years, standard deviation of 2.5 years) gave informed written consent to participate for monetary compensation ($10 per hour). All participants were right-handed, had normal hearing and normal or corrected-to-normal vision, and had no history of concussion. They were tested individually in a dimly lit or darkened room in the period between 5/11/2017 and 1/27/2020. The study protocol was approved by the Northwestern University Institutional Review Board. A group of twenty-four individuals participated in a rest-with-eyes-closed condition in which EEG was recorded for ∽5 min while participants rested with their eyes closed and freely engaged in spontaneous thoughts. A second (non-overlapping) group of twenty-four individuals participated in a replication of the rest-with-eyes-closed condition and subsequently participated in a rest-with-eyes-open-in-the-dark condition which was the same as the former except that the room was darkened and participants kept their eyes open while blinking naturally. A third group of twenty-one individuals (seventeen of whom previously participated in the rest-with-eyes-closed condition) participated in a silent-nature-video condition in which EEG was recorded for ∽5 min while they viewed a silent nature video. To evaluate test-retest reliability, the silent-nature-video condition was run twice (20-30 min apart), referred to as earlier viewing and later viewing. A generic nature video was presented on a 13-inch, 2017 MacBook Pro equipped with 2880(H)-by-1800(V)-pixel-resolution LCD display with normal brightness and contrast settings, placed 100 cm in front of participants, subtending approximately 16° (H)-by-10° (V) of visual angle. Subsets of these data were previously analyzed for different purposes (Menceloglu et al., 2020a, 2020b, 2021a, 2021b, 2021c).

### EEG recording and pre-processing

EEG was recorded from 64 scalp electrodes (although we used a 64-electrode montage, we excluded signals from noise-prone electrodes, Fpz, Iz, T9, and T10, from analyses) at a sampling rate of 512 Hz using BioSemi ActiveTwo system (see www.biosemi.com for details), while participants rested with their eyes closed, rested with their eyes open in the dark, or viewed a silent nature video for approximately 5 min. Electrooculographic (EOG) activity was monitored using four face electrodes, one placed lateral to each eye and one placed beneath each eye. Two additional electrodes were placed on the left and right mastoids. The EEG data were preprocessed using EEGLAB and ERPLAB toolboxes for MATLAB (Delorme and Makeig, 2004; Lopez-Calderon and Luck, 2014). The data were re-referenced offline to the average of the two mastoid electrodes and high-pass filtered at 0.01 Hz to remove drifts.

### Estimating dura sources by surface-Laplacian transforming EEG signals

EEG source reconstruction methods constrained by structural MRI and fMRI localizers obtained from each participant can achieve superior source reconstruction with a brain model customized for each participant (Cottereau et al., 2015). Such approach, however, was unavailable to us as we had neither structural MRI nor fMRI data for our participants. Among the non-customized source-imaging methods, we chose the surface-Laplacian transform that (theoretically) estimates the spatial distribution of macroscopic current sources and sinks on the dura surface. The surface-Laplacian transform has been shown to produce similar sources to those inferred by deconvolving scalp EEG potentials using a generic model of thicknesses and impedances of scalp and skull (Nunez et al., 1994). The surface-Laplacian transform has also been shown to approximate simulated sources and/or to extract neural correlates of some behavioral performances to a similar degree to commonly used source-imaging methods such as sLORETA and Beamforming (Cohen, 2015; Tenke and Kayser, 2012). Further, there is no evidence (to our knowledge) to suggest that the latter source-imaging methods provide greater spatial resolution than the surface-Laplacian transform. Thus, our preference was to use the surface-Laplacian transform which is the most general source-imaging method that relies the least on model-specific assumptions and free parameters (Hjorth, 1980; Nunez et al., 1994; Nunez and Srinivasan, 2006; Kayser and Tenke, 2006, 2012).

The surface-Laplacian transform is expected to reduce volume-conduction effects from substantially greater than 5 cm in raw EEG to within 1-3 cm (Cohen, 2014; Menceloglu et al., 2021c; Nunez et al., 1994; Tenke and Kayser, 2012) which approximately corresponded to the average spacing of electrodes in our 64-channel montage. For our implementation of the surface-Laplacian transform, we used Perrin and colleagues’ algorithm (Perrin et al., 1987, 1989a, 1989b) with a “smoothness” value, λ = 10^−5^ (recommended for 64 channels; Cohen, 2014). We refer to the surface-Laplacian transformed EEG signals that represent the macroscopic current sources and sinks on the dura surface under the 60 scalp sites (with the four noise-prone sites excluded from the analyses) simply as EEG signals.

### EEG analysis

#### Taking the temporal derivative

Our goal was to track spectral maxima (likely indicative of oscillation frequencies) at sub-second temporal resolution. The general 1/*f*^*β*^ spectral background in EEG interferes with the identification of spectral maxima. A commonly employed strategy to circumvent this problem is to compute a Fourier transform over partially overlapping time windows of several seconds or longer, and then fit a power function to each time-windowed Fourier transform to subtract out the 1/*f*^*β*^ component (e.g., van Albada and Robinson, 2013; Chiang et al., 2011; Donoghue et al., 2020). However, it is difficult to reliably estimate the 1/*f*^*β*^ component on a sub-second timescale. Note that *β* in the 1/*f*^*β*^ spectral background varies around 1 (see He (2014) for a review of the various factors that influence *β*, and Gao et al. (2017) for contributions of the excitatory and inhibitory neural dynamics to *β*). Although taking the temporal derivative of EEG (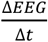, where Δ*t* is the temporal resolution, i.e., 1/512 sec here) would completely neutralize the 1/*f*^*β*^ component only when *β* = 1, the method worked well for our data (Menceloglu et al., 2020a, 2021a, 2021b, 2021c), suggesting that *β*∽1 generally held for our data. As we discussed in our prior reports (Menceloglu et al., 2021a, 2021b, 2021c), taking EEG temporal derivative offers additional advantages. For example, EEG temporal derivatives are drift free. EEG temporal derivatives may be considered a deeper measure of neural activity than EEG in the sense that scalp-recorded potentials are generated by the underlying neural currents and taking EEG temporal derivative macroscopically estimates those currents (as currents in an RC circuit are proportional to the temporal derivative of the corresponding potentials). Further, there is some evidence that EEG temporal derivatives offer a more effective neural measure than EEG for brain-computer interface (Andreou and Poli, 2016).

#### Time-frequency decomposition using Morlet wavelets

To track EEG power spectra at high temporal and spectral resolutions, we used a Morlet wavelet-convolution method suitable for time-frequency decomposition of signals containing multiple oscillatory sources of different frequencies (see Cohen (2014) for a review of different methods for time-frequency decomposition). Each Morlet wavelet is a Gaussian-windowed complex sinusoidal template characterized by its frequency as well as its temporal and spectral widths that limit its temporal and spectral resolutions. We convolved each EEG waveform (i.e., its temporal derivative) with a set of wavelets tuned to a range of frequencies, yielding a time series of complex values per wavelet frequency. The power and phase of each extracted sinusoidal component at each time point were given by the modulus squared (power) and the arc tangent of the ratio of the imaginary component to the real component (phase). We used a set of wavelets with 160 frequencies, *f*_*w*_’s, ranging from 5 Hz to 15 Hz. The *f*_*w*_’s were logarithmically spaced as neural temporal-frequency tunings tend to be approximately logarithmically scaled (e.g., Hess and Snowden, 1992; Lui et al., 2007). The accompanying *n* factor (roughly the number of cycles per wavelet, with the precise definition, *n* = 2*πf* · *SD*, where *SD* is the wavelet standard deviation) was also logarithmically spaced between 11.7 and 35. This spacing yielded a temporal resolution of *SD* = 370 ms and a spectral resolution of, *FWHM* (*full width at half maximum of wavelet spectrum*) = 1.0 Hz, that were virtually invariant across the wavelet frequencies.

#### Identifying oscillation frequencies per time point

When multiple oscillations at neighboring frequencies coincide, each oscillation frequency does not necessarily generate a local maximum in the power spectrum due to a limited spectral resolution. An example is shown in Figure 1A, plotting a single-time-point power spectrum obtained with our Morlet wavelets for a simulated data containing five sinusoidal oscillations at 6, 7, 9.5, 10.5, and 13 Hz (vertical arrows) of different amplitudes. Oscillation frequencies are typically identified as spectral peaks that theoretically coincide with points of phase stationarity (e.g., Mallat, 2009). However, only three of the five oscillation frequencies were identified as spectral peaks (black open circles in Figure 1A) which coincided with points of phase stationarity (negatively sloped zero crossings of 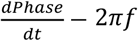 shown in Figure 1B). Spectral peaks and phase stationarity missed the low-amplitude oscillations at 7 Hz and 9.5 Hz in the neighborhoods of the high-amplitude oscillations at 6 Hz and 10.5 Hz (Figure 1A).

**Figure 1.**
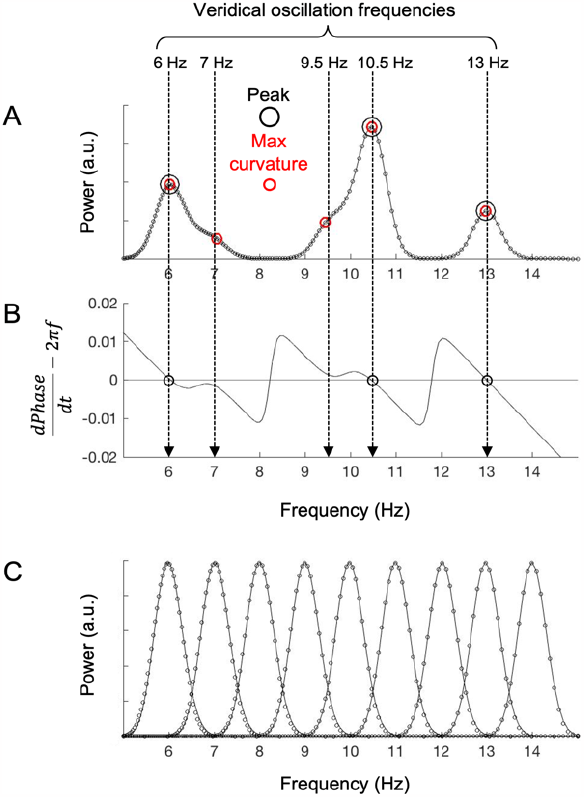
Identifying oscillation frequencies as curvature maxima in a power spectrum (per time point). **A**. A power spectrum of simulated EEG data (at a single time point) containing sinusoidal oscillations at 6, 7, 9.5, 10.5, and 13 Hz (vertical arrows) of various amplitudes, processed with our 160 wavelets (see text). The large black open circles indicate oscillation frequencies identified as spectral peaks, whereas the small red open circles indicate oscillation frequencies identified as curvature maxima. Note that only curvature maxima identified the low-amplitude oscillations at 7 Hz and 9.5 Hz occurring near the high-amplitude oscillations at 6 Hz and 10.5 Hz. **B**. A plot of 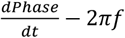 as a function of *f* (frequency). The negatively sloped zero crossings (black open circles) indicate the points of phase stationarity, which coincide with the spectral peaks. **C**. Representative point-spread functions (power spectra of single sinusoidal oscillations) showing the spectral resolution of our 160-wavlet spectral decomposition. The point-spread functions are well fit by Gaussian functions (continuous curves).

We thus used a more sensitive method to identify oscillation frequencies. As shown in Figure 1C, our wavelet-based time-frequency decomposition generated Gaussian-shaped power spectra for single-frequency oscillations (point-spread functions). We took advantage of the fact that the curvature of a Gaussian function (maximum at its peak) falls faster than its value. Thus, local curvature maxima provide a higher spectral resolution for identifying oscillation frequencies than spectral peaks. For example, the low-amplitude oscillations at 7 Hz and 9.5 Hz generated curvature maxima (red open circles in Figure 1A) though they did not generate spectral peaks (or points of phase stationarity). We tracked curvature maxima as a function of time at anterior-posterior sites along the midline axis (AFz, Fz, FCz, Cz, CPz, Pz, POz, and Oz) as well as along the left-hemisphere axis (AF3, F3, FC3, C3, CP3, P3, PO3, and O1) and the right-hemisphere axis (AF4, F4, FC4, C4, CP4, P4, PO4, and O2).

The number of identified curvature maxima per time point was consistent across participants and behavioral conditions, showing nearly identical distributions centered around eight (Figure 2). It is likely that some of these curvature maxima were due to noise. Nevertheless, instead of imposing an arbitrary power threshold, we analyzed data relative to their time-shuffled controls as described below. Spurious effects due to noise would be statistically equivalent for the actual data and their time-shuffled controls.

**Figure 2.**
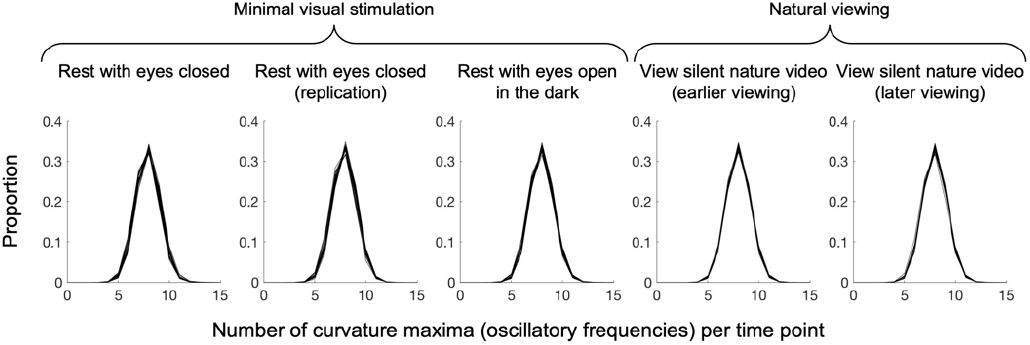
Distribution of the number of curvature maxima (oscillation frequencies) per time point for the five behavioral conditions. The distributions were averaged across all sites as they did not appreciably differ across sites. Each curve represents the distribution from a single participant in the corresponding condition. Note that the distributions were similar across all participants and behavioral conditions.

## Results

Our goal was to uncover general rules governing spontaneous oscillation-frequency convergences across anterior-posterior sites along the midline, left-hemisphere, and right-hemisphere axes. Extensive oscillation-frequency convergences involving nearly all eight anterior-posterior sites were rare when we required exact alignments of oscillation frequencies across sites. We thus registered oscillation-frequency convergences with a tolerance commensurate with the spacing of our 160 wavelet frequencies. Specifically, we broadened each identified oscillation frequency at each site (at each time point) by ±1 unit of our wavelet-frequency spacing, which logarithmically increased from ±0.04 Hz at 5 Hz to ±0.11 Hz at 15 Hz due to our logarithmic spacing of the 160 wavelet frequencies (see Materials and methods). That is, we considered oscillation frequencies to be convergent when they were within ±0.04 Hz at the lowest end and within ±0.11 Hz at the highest end.

For each site (for each participant), we represented the temporal flows of oscillation frequencies in an oscillation-frequency-(row)-by-time-(column) matrix—the *flow matrix*.

To implement the above-mentioned tolerance for frequency-convergence detection, each identified oscillation frequency at each time point as well as the frequency values immediately above and below it were given the value of 1, with the rest of the matrix filled with zeros. The temporal flows of spontaneous oscillation-frequency convergences were then obtained by stacking the flow matrices from the eight anterior-posterior sites on top of one another and summing them across the vertical dimension, yielding an oscillation-frequency-(row)-by-time-(column) matrix where values one through eight represented oscillation-frequency convergences across one through eight sites. We call this the *flow-convergence matrix*.

How oscillation frequencies dynamically converged across anterior-posterior sites can be visualized by plotting a flow-convergence matrix. An example is shown in Figure 3 for one participant in the rest-with-eyes-closed condition where cooler/darker colors indicate frequency convergences across larger numbers of midline anterior-posterior sites. Our goal was to uncover general characteristics of these spontaneous oscillation-frequency convergences to infer their mechanisms and functions.

**Figure 3.**
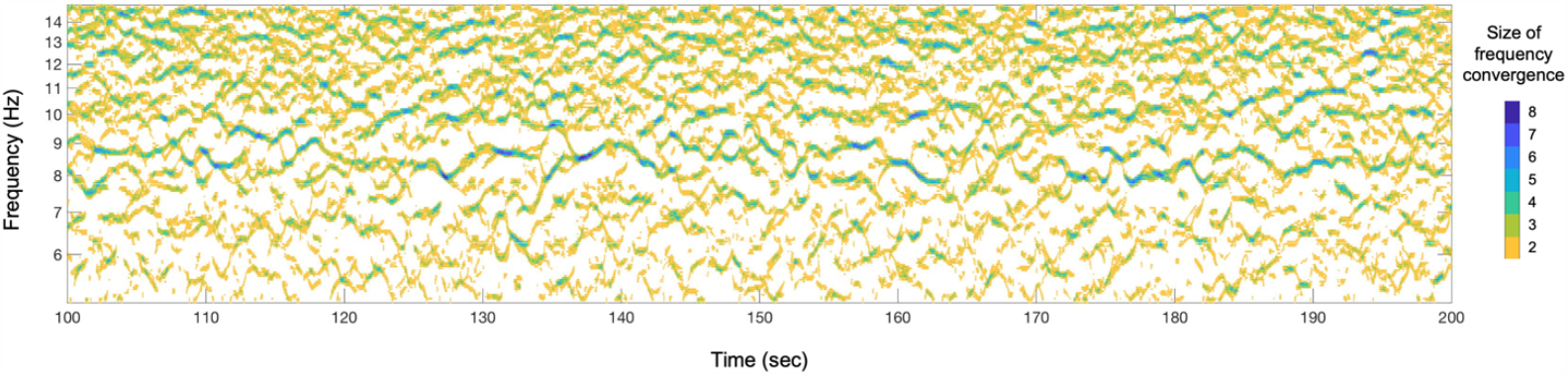
Visualization of a flow-convergence matrix, showing the number of frequency-converged sites (color coded) along the midline anterior-posterior axis as a function of oscillation frequency (y-axis) and time (x-axis), for one participant in the rest-with-eyes-closed condition during a 100-sec period. Cooler/darker colors indicate oscillation-frequency convergences across larger numbers of anterior-posterior sites.

We first verified that spontaneous oscillation-frequency convergences were driven. As oscillations in the extended alpha range may contain dominant frequencies (e.g., ∽10 Hz peaks), spontaneous frequency convergences may simply reflect coincidences of dominant oscillations across sites. We ruled out this possibility by comparing the total durations of oscillation-frequency convergences of one through eight sites between the actual flow-convergence matrices and their time-shuffled controls. We generated a time-shuffled version by randomly time-shifting the flow matrices of each site by up to 150 sec (a half of the total duration of ∽5 min) prior to summing across sites. If the instances of oscillation-frequency convergences were due to coincidences of dominant oscillations, the total durations of frequency convergences should be equivalent for the actual and time-shuffled flow-convergence matrices. Total durations of frequency convergences across one through eight sites can be approximately computed by tallying the total instances of 1’s through 8’s in a flow-convergence matrix. These values overestimated the actual total durations by up to three-fold as we broadened each oscillation-frequency flow by ±1 wavelet-frequency spacing to increase the tolerance of convergence detection (see above). Nevertheless, this was not a problem as we compared the actual total durations with those averaged from 20 time-shuffled controls (per behavioral condition per participant) given that we computed total durations of convergences across one through eight sites in the same way for the actual and time-shuffled flow-convergence matrices.

As shown in Figure 4, the total durations of convergences of four or more sites were consistently elevated in the actual data (red curves) relative to the time-shuffled controls (black curves) for all participants in all behavioral conditions. Given that the total durations are plotted in log scale, the elevations were substantial.

**Figure 4.**
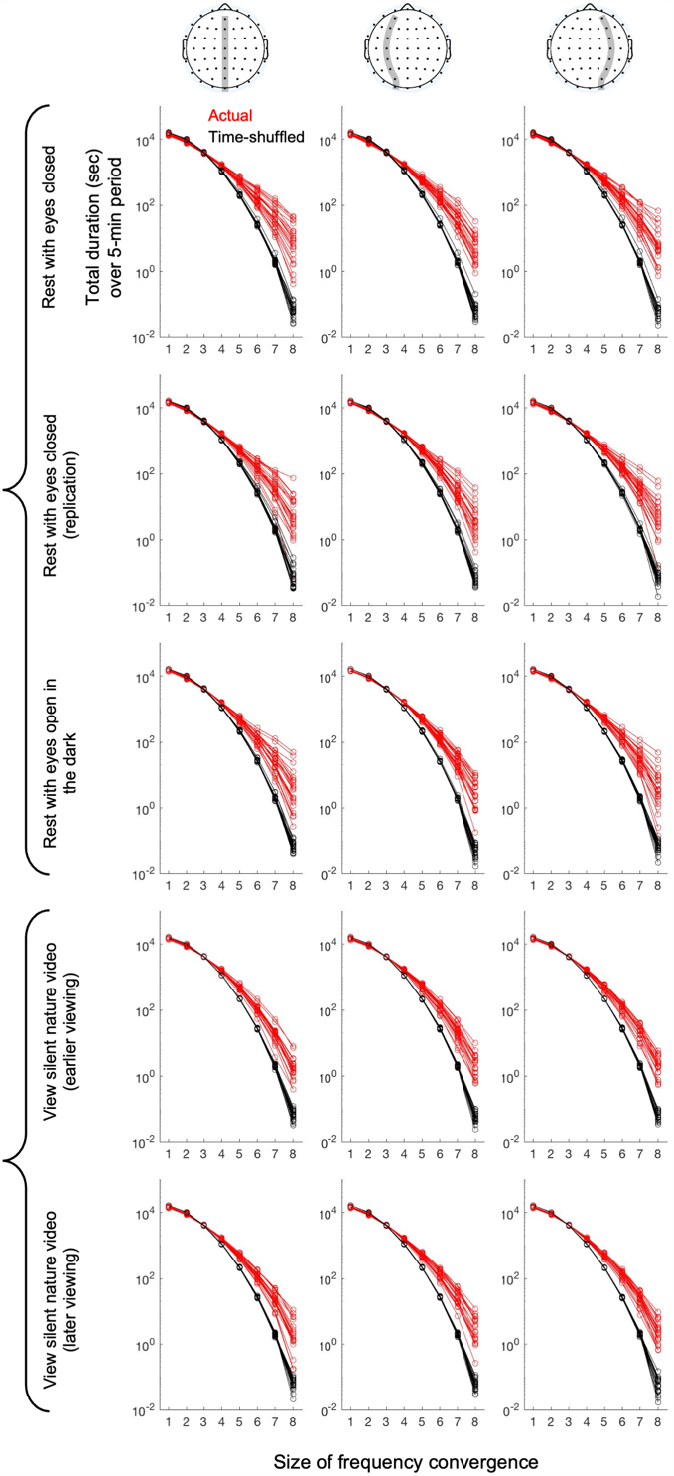
Approximate (see text) total duration (sec) of anterior-posterior oscillation-frequency convergence as a function of the size of frequency convergence, along the midline axis (first column), the left-hemisphere axis (second column), and right-hemisphere axis (third column), for the five behavioral conditions (rows). The actual data are shown in red and the time-shuffled controls (averaged across 20 versions) are shown in black. Each curve represents a participant. Note that the durations of frequency convergences of four or more sites were elevated in the actual data (red) relative to the time-shuffled controls (black) for all participants in all conditions.

Having verified that oscillation-frequency convergences were likely driven (rather than coincidences), we examined general characteristics of anterior-posterior oscillation-frequency convergences. While a variety of spatial, spectral, and temporal features of oscillation-frequency convergences can be investigated, we primarily focused on the following three questions. First, what drives large oscillation-frequency convergences? Second, how are oscillation-frequency convergences related to oscillatory power? Third, do oscillation-frequency convergences generate consistent anterior-posterior phase relations? As discussed below, investigating these questions provided some insights into a potential mechanism and function of anterior-posterior oscillation-frequency convergences.

What mechanism may facilitate the spreading of anterior-posterior oscillation-frequency convergences? We reasoned that if synergistic interactions drove oscillation-frequency convergences, the propensity to converge would become stronger as more sites converged. We assessed this possibility by computing the probability of growing in convergence as a function of the number of converged sites. Using the flow matrix for each site (containing 1’s where oscillations were identified, broadened by ±1 wavelet-frequency spacing; see above), we computed the average probability of frequency convergence across *k* sites. We first obtained the average base probability of oscillatory activity by computing the proportion of 1’s in the flow matrix of each site and averaging the values across all sites. We call this *P*(1), the average probability of oscillatory activity. If oscillation-frequency convergences were due to coincidences, the average probability of frequency convergences across *k* sites, *P*(*k*), should be given by *P*(1)^*k*^.

We computed the probability of frequency convergences across *k* > 1 sites as follows. For each combination of *k* sites, we stacked and summed their flow matrices and then computed the proportion of *k*’s in the resultant matrix to obtain the probability of *k* convergence. We then averaged the probabilities computed for all combinations of *k* sites to obtain the average probability of *k* convergence, *P*(*k*). To estimate the probability of growing in frequency convergence at each level of convergence (*k*), we computed the conditional probability of obtaining *k* convergence given *k*– 1 convergence, *P*(*k*|*k*– 1), by dividing *P*(*k*) by *P*(*k*– 1) because *k* convergence was a subset of *k*– 1 convergence. We computed *P*(*k*|*k*– 1) (where 2 ≤ *k* ≤ 8) for each participant for each behavioral condition.

As shown in Figure 5A, in all five behavioral conditions (rest with eyes closed, its replication, rest with eyes open in the dark, and the two instances of viewing a silent nature video) the probability of growing in convergence, *P*(*k*|*k*– 1), substantially increased as a function of the number of frequency-converged sites, *k* (red curves). Slight increases were also observed in the time-shuffled controls (based on averaging 20 time-shuffled data) (black curves), indicating that some features of the flow matrices such as major oscillation flows occurring in relatively narrow frequency ranges slightly increased the probability of coincidental convergences relative to the chance level (gray lines). The data for large frequency convergences (for *k* = 7 and especially for *k* = 8) were somewhat noisy as their instances were relatively rare especially for the time-shuffled controls (Figure 4). Importantly, *P*(*k*|*k*– 1) for the actual data generally increased more steeply than for the time-shuffled controls as a function of *k* (Figure 5B). This indicates that the probability of growing in frequency convergence increased as a function of the size of convergence, controlling for any potential effects of time independent spectral features of the flow matrices at the individual sites that might have increased the probability of coincidental frequency convergences. We thus conclude that synergistic interactions drove oscillation-frequency convergences across anterior-posterior sites.

**Figure 5.**
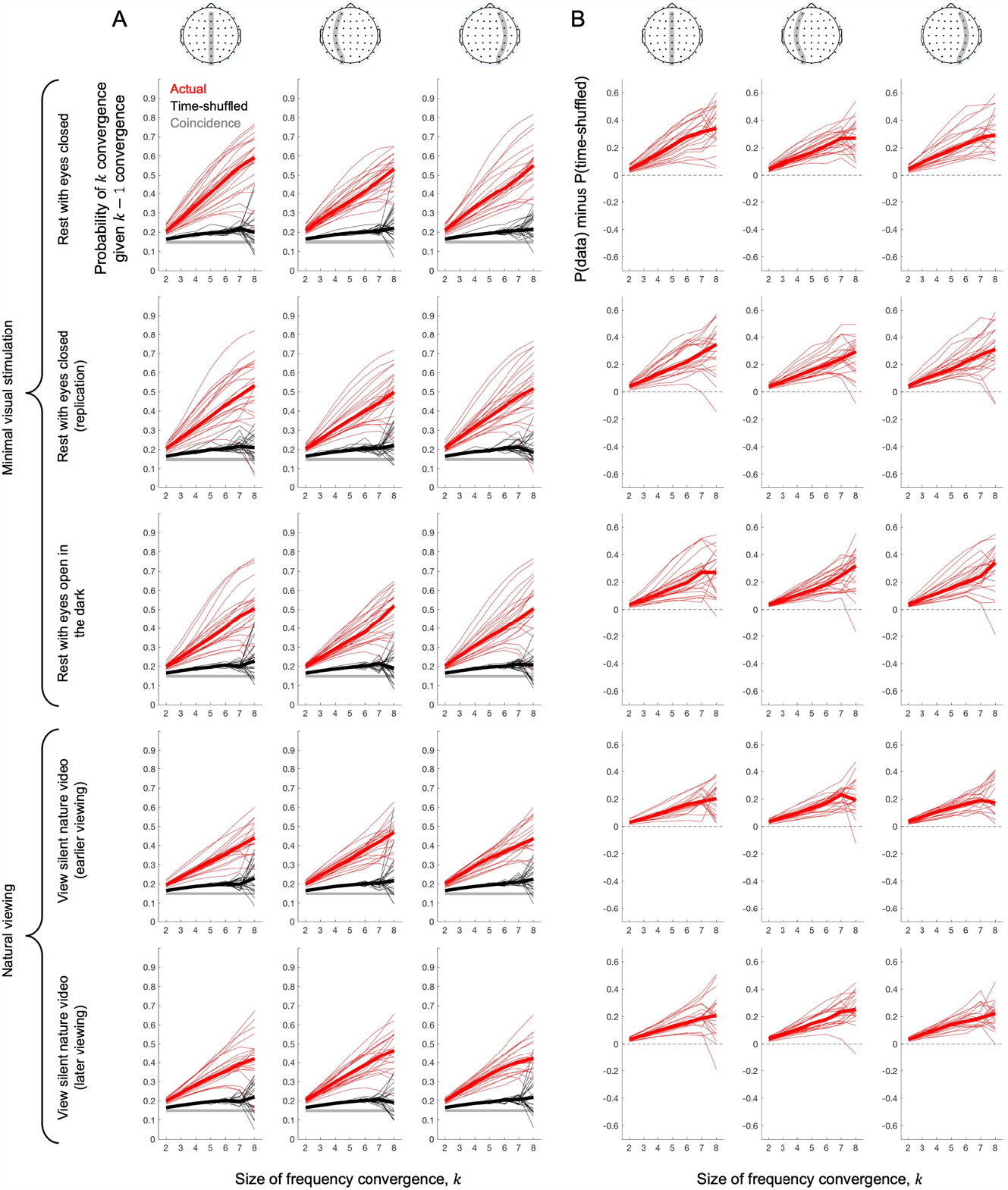
The probability of *k* convergence (oscillation-frequency convergence across *k* sites) given *k* − 1 convergence, that is, the probability of growing in convergence from *k* − 1 to *k, P*(*k*|*k*– 1), as a function of *k* for the five behavioral conditions (rows). **A**. *P*(*k*|*k*– 1) for the actual data (red), their time-shuffled controls (black, averaged across 20 versions), and the probabilities expected if oscillation-frequency convergences occurred randomly (gray). **B**. *P*(*k*|*k*– 1) for the actual data minus *P*(*k*|*k*– 1) for the corresponding time-shuffled controls. In both parts A and B, the thin lines show individual participants’ values while the thick lines show the median values. The data are somewhat noisy for high convergences (for *k* = 7 and especially for *k* = 8) as their instances were relatively rare especially for the time-shuffled controls (see Figure 4). The data for the anterior-posterior sites along the midline axis, the left-hemisphere axis, and the right-hemisphere axis are shown in the first, second, and third column, respectively. Note that the probability of increasing in frequency convergence generally increased as the number of frequency-converged sites increased, suggesting that synergistic interactions drive oscillation-frequency convergences across anterior-posterior sites.

What mechanism might mediate the synergistic interactions? Oscillatory neural synchronizations occurring in multiple regions at a matched frequency may enhance one another (e.g., Moon et al., 2015; Tewarie et al., 2019). In turn, the increased oscillatory synchronizations in those frequency-matched regions may facilitate the spreading of oscillatory synchronizations to neighboring regions. In our EEG data, regional oscillatory synchronizations are reflected in wavelet-extracted oscillatory power. Thus, if synergistic oscillation-frequency convergences were mediated by convergence-driven inter-regional enhancements of oscillatory synchronizations, larger frequency convergences should be associated with higher oscillatory power at the participating sites.

To evaluate this possibility, we temporally averaged oscillatory power at each site separately for each size of frequency convergence, *k*. For instance, we averaged oscillatory powers at AFz when its oscillation frequencies did not converge with any other sites (*k* = 1), when its oscillation frequencies converged with one other site (*k* = 2), when its oscillation frequencies converged with two other sites (*k* = 3), and so on. This yielded the average oscillatory power at each site as a function of the size of frequency convergence in which it participated. However, we may obtain a spurious association between higher power and larger frequency convergence due to general frequency-flow characteristics. For example, if higher-power oscillations tended to be concentrated within a narrow frequency range at individual sites while lower-power oscillations were broadly distributed across frequencies, higher-power oscillations would coincide with higher probability than lower-power oscillations. To account for such confounds, we applied the same analysis to the corresponding time-shuffled controls. Any biased coincidences due to general frequency-flow characteristics would be equivalent for the actual data and their time-shuffled controls. Thus, for each site, we computed its average oscillatory power minus the value obtained with the corresponding time-shuffled control (averaged across 20 versions) as a function of the size of frequency convergence in which it participated. To facilitate comparisons of these power-by-convergence functions across sites, behavioral conditions, and participants, we normalized each function (actual minus control) by dividing it by its maximum actual power.

Figure 6 shows the (normalized) average oscillatory power at each site (color coded from dark blue [posterior] to yellow [anterior]) as a function of the size of frequency convergence in which it participated. The open circles indicate Bonferroni-corrected (for eight convergence sizes) significant difference from the control level (i.e., zero) at α=0.05 (two-tailed). In all five behavioral conditions, the oscillatory power generally increased as a function of the size of frequency convergence at all sites (except at the anterior-most site in some cases; see yellow curves in Figure 6), rising significantly above the control levels at a convergence of four or five sites and steeply increasing at larger convergences. These increases were somewhat steeper under minimal visual stimulation than in natural viewing (the top three vs. bottom two rows in Figure 6). Overall, the results summarized in Figure 6 are consistent with the aforementioned possibility that synergistic oscillation-frequency convergences may be mediated by inter-regional enhancements of oscillatory synchronization.

**Figure 6.**
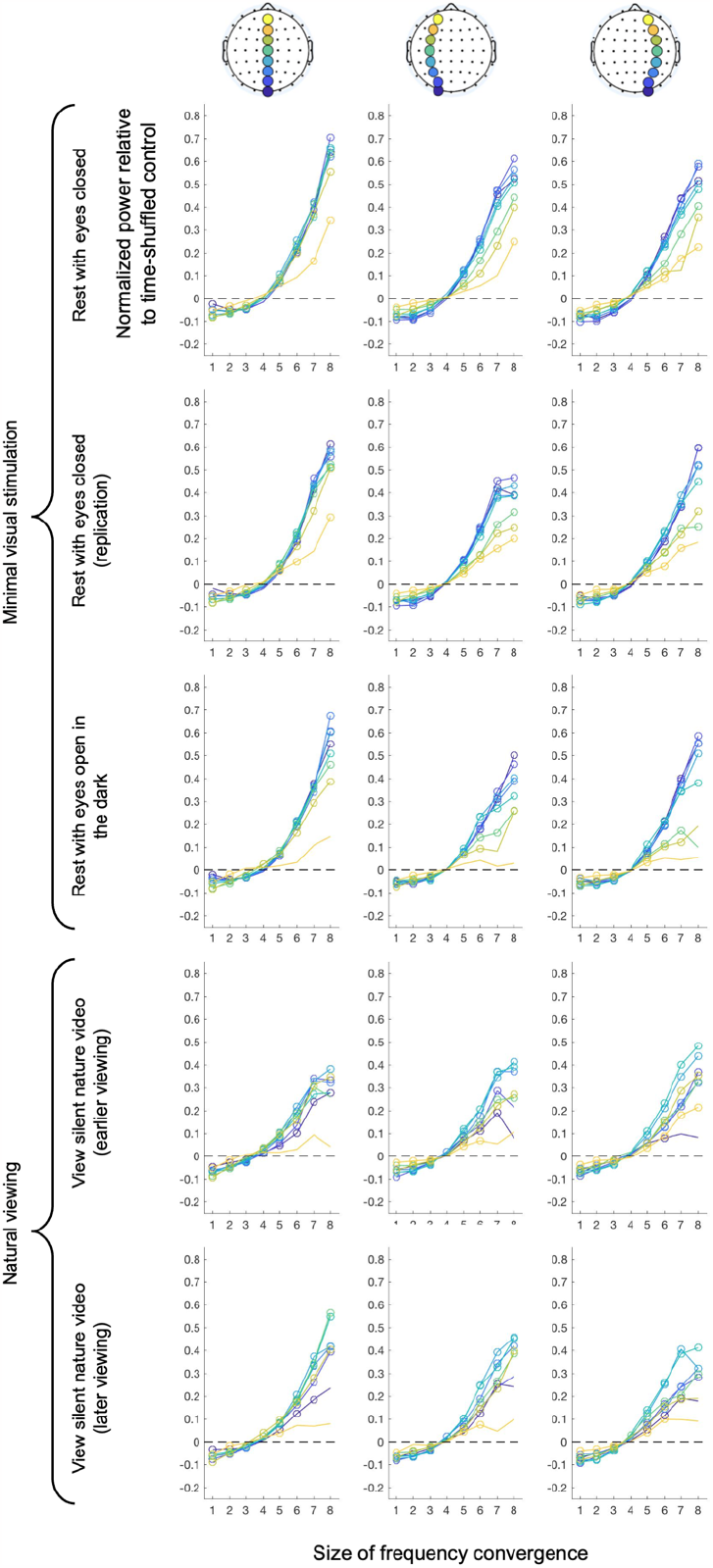
Time-averaged oscillatory power at anterior-posterior sites (color-coded from dark blue [posterior] to yellow [anterior]) as a function of the size of frequency convergence, along the midline axis (first column), left-hemisphere axis (second column), and right-hemisphere axis (third column) for the five behavioral conditions (rows). The power values were computed relative to those obtained with the corresponding time-shuffled controls (averaged across 20 versions) and normalized to the maximum actual power for each function (per site per behavioral condition per participant). The values averaged across participants are plotted, with each open circle indicating Bonferroni-corrected (for eight convergence sizes) statistical significance (*α* = 0.05, two-tailed) relative to zero (control levels). Note that most sites showed accelerated increases in power when they participated in frequency convergences of four or more sites.

To gain some insights into how anterior-posterior frequency convergences spread, we examined each site’s propensities for joining frequency convergences of different sizes. For example, in a simple case where frequency convergences always spread sequentially from site 1 to site 8, frequency convergences of two sites would predominantly involve sites 1 and 2, convergences of three sites would predominantly involve sites 1, 2, and 3, convergences of four sites would predominantly involve sites 1, 2, 3, and 4, and so on. Propensities for each site to be included in frequency convergences of different sizes *k* were computed as the ratio of the actual instances to those obtained with the corresponding time-shuffled controls (averaged across 20 time-shuffles). For example, a value of 0.8 would indicate that the number of instances of a particular site being involved in frequency convergences of size *k* was 80% relative to the corresponding control level, and a value of 3.5 would indicate that the number of instances was 3.5 times as frequent as the corresponding control level, with 1 indicating the level expected from coincidences given the time independent spectral features of the flow matrices at the individual sites.

Anterior-posterior oscillation-frequency convergences appear to spread differently along the midline axis than the left- and right-hemisphere axes. Along the midline axis, the instances of participating in frequency convergences of three to seven sites were consistently reduced for the posterior-most and anterior-most sites. Note that the instances of participating in frequency convergences of all eight sites were necessarily equal for all sites. The flattened inverted-U pattern was consistently observed in all behavioral conditions (black curves in Figure 7A) and for most participants (gray curves in Figure 7A). For frequency convergences of only two sites and singletons, we obtained the pattern that was opposite to those in

**Figure 7.**
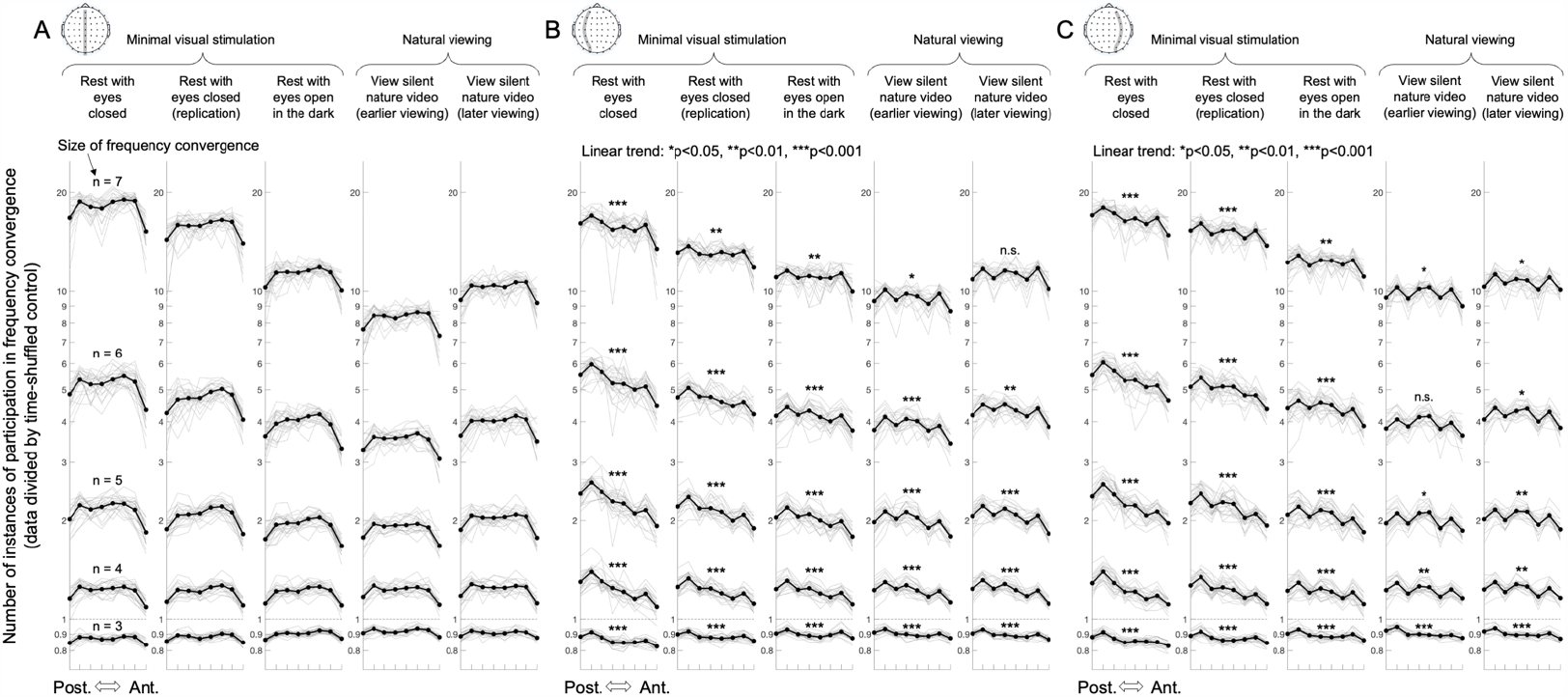
The number of instances of each site along the anterior-posterior axis participating in frequency convergences of three to seven sites, along the midline (**A**) and in the left (**B**) and right (**C**) hemispheres, for the five behavioral conditions (columns). The values were divided by the corresponding values obtained with the time-shuffled controls (averaged across 20 versions). For example, a value of 0.8 indicates that the actual instances were 80% of the control instances, a value of 3.5 indicates that the actual instances were 3.5 times as frequent as the control instances, with 1 indicating the level expected from coincidences given the time independent spectral features of the flow matrices at the individual sites. The solid curves show the geometric means while the gray curves show the values from individual participants aligned at the corresponding geometric means. Note that along the midline axis (A), the posterior-most and anterior-most sites were less likely to be included in frequency convergences of three or more sites, whereas along the left-hemisphere (B) and right-hemisphere (C) axes, more anterior sites were less likely to be included in frequency convergences of three or more sites, yielding significant negative slopes.

Figure 7A (not shown), indicating that the posterior-most and anterior-most sites had elevated tendencies to generate isolated oscillations that did not spread beyond a pair of sites. Overall, these results suggest that frequency convergences along the midline axis can develop from most sites except that the posterior-most and anterior-most sites tend to join later.

Along the left- and right-hemisphere axes, the instances of participating in frequency convergences of three to seven sites were progressively diminished from posterior to anterior sites, with the negative monotonicity (computed as Fisher Z transformed Spearman’s *r*) being statistically significant in most cases across all behavioral conditions (Figure 7B and 7C). It appears that the negative monotonicity was generally stronger under minimal visual stimulation than in natural viewing. As above, these patterns reversed for frequency convergences of only two sites and singletons (not shown), indicating that the sites that participated less in larger frequency convergences had elevated tendencies to generate isolated oscillations that did not spread beyond a pair of sites. Overall, these results suggest that along the left -and right-hemisphere axes frequency convergences tend to spread from posterior to anterior sites.

Note that the instances of participating in frequency convergences of three sites were below the control levels in all cases (Figure 7A, 7B, and 7C), suggesting that synergistic frequency convergences operate on a threshold of more than three converged sites. This is consistent with the results above that oscillatory power was also consistently below the control levels for frequency convergences of three or less sites (Figure 6).

Given that anterior-posterior oscillation-frequency convergences appear to be synergistically driven, what functions might they serve? Investigating this question would require explorations of behavioral tasks whose performances are influenced by intrinsic oscillation-frequency convergences. Here we indirectly investigated potential functions of oscillation-frequency convergences by examining the phase relations induced by frequency convergences. For example, if frequency convergences induced a phase gradient with progressive phase lags from anterior to posterior sites, one could infer that frequency convergences facilitate a flow of information from anterior to posterior regions.

We computed sequential phase differences from posterior to anterior. For example, for the anterior-posterior axis along the midline, we computed the phase of POz minus the phase of Oz, the phase of Pz minus the phase of POz, the phase of CPz minus the phase of Pz, the phase of Cz minus the phase of CPz, the phase of FCz minus the phase of Cz, the phase of Fz minus the phase of FCz, and the phase of AFz minus the phase of Fz. These sequential phase differences were computed wherever the size of frequency convergence was two or larger in the flow-convergence matrices (e.g., Figure 3), and were averaged (as complex angles) across all instances but separately for each size of frequency convergence. Because of the temporal averaging, this analysis detected phase gradients only if they were consistent over the ∽5 min period.

To visualize the phase gradients, we spatially integrated the sequential pairwise phase differences from the posterior to anterior sites while arbitrarily setting the phase at the posterior-most site to zero (i.e., we cumulatively summed the sequential pairwise phase differences from posterior to anterior). A negative phase gradient (a negative slope) indicates that anterior sites increasingly lagged in phase, suggestive of a flow of information in the anterior direction, whereas a positive phase gradient (a positive slope) indicates that posterior sites increasingly lagged in phase, suggestive of a flow of information in the posterior direction.

Upon examination of the phase gradients obtained from each participant, we noted that they classified into two prototypes. Clear examples of these prototypes are shown in Figure 8B. One prototype consisted of generally negative phase gradients from posterior to anterior (e.g., Figure 8B, left), whereas the other prototype consisted of generally positive phase gradients from posterior to anterior (e.g., Figure 8B, middle and right). Both gradients became steeper as the size of frequency convergence increased.

**Figure 8.**
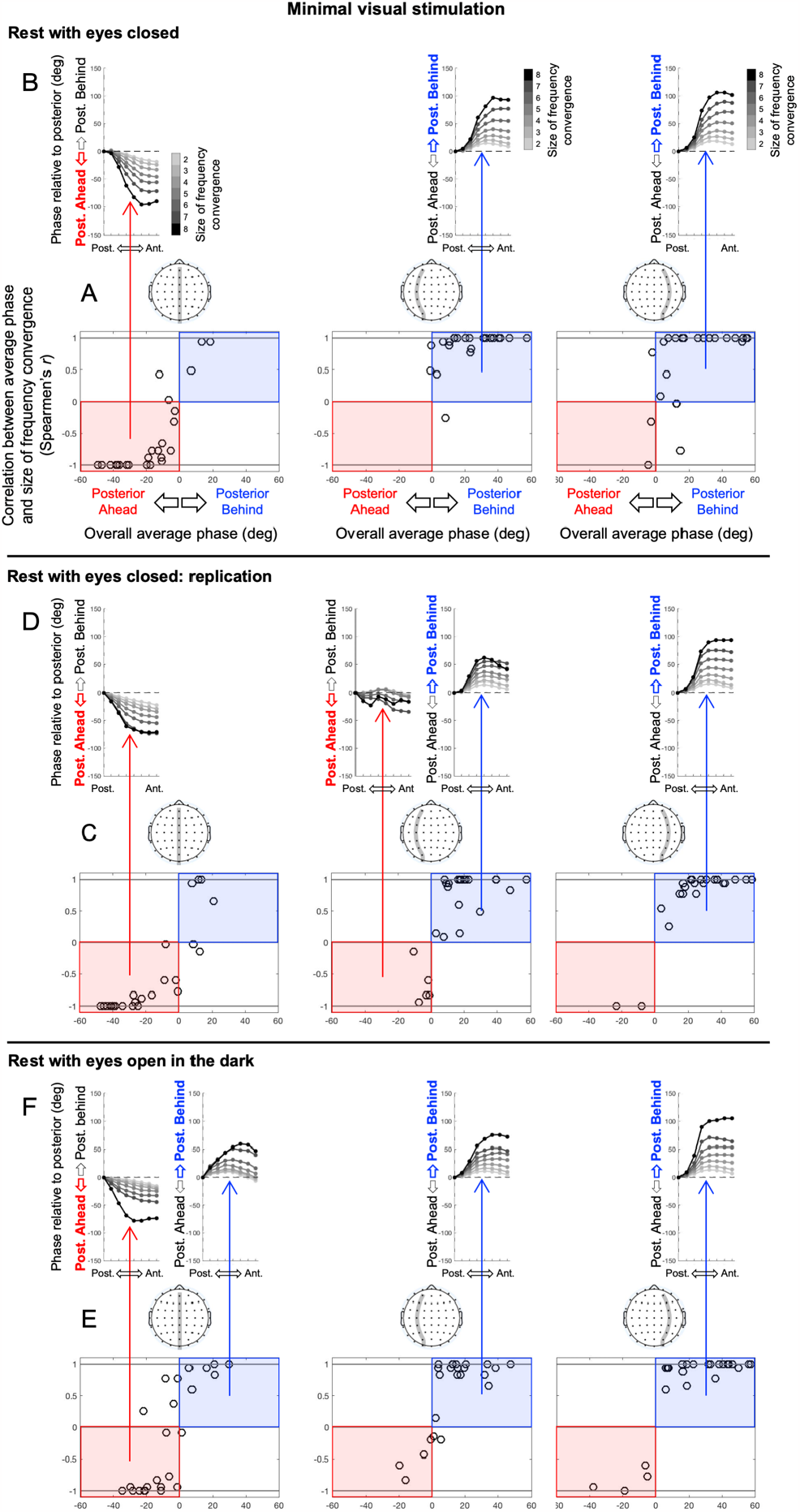
Two general types of posterior-to-anterior phase gradients observed during inter-site oscillation-frequency convergences under minimal visual stimulation: the rest-with-eyes-closed (A and B), its replication (C and D), and the rest-with-eyes-open-in-the-dark (E and F) conditions. **ACE**. Distribution of participants in the two-dimensional space defined by the overall average phase (averaged across frequency-convergence sizes and anterior-posterior sites) on the x-axis and the correlation between the average phase (averaged across anterior-posterior sites) and the size of frequency convergence (Spearmen’s *r*) on the y-axis, shown separately for the three anterior-posterior axes: the midline axis (left panel), the left-hemisphere axis (middle panel), and the right-hemisphere axis (right panel). The x=0 and y=0 lines divide the two-dimensional space into four quadrants with each circle representing a participant. Note that most participants populated the lower-left quadrant for the midline axis (red rectangles in the left column) whereas most participants populated the upper-right quadrant for the left- and right-hemisphere axes (blue rectangles in the middle and right columns). **BDF**. Average phase gradients for all oscillation-frequency convergence sizes (gray-scale coded) shown for each quadrant that included five or more participants (averaged across relevant participants). Note that the lower-left quadrant is characterized by generally negative posterior-to-anterior phase gradients that became steeper (more negative) as the size of frequency convergence increased. In contrast, the upper-right quadrant is characterized by generally positive posterior-to-anterior phase gradients that became steeper (more positive) as the size of frequency convergence increased.

To determine how consistently the phase-gradient patterns from individual participants classified into these prototypes, we examined the distribution of phase-gradient patterns in the two-dimensional space defined by (1) the overall average phase and (2) the correlation between the average phase and the size of frequency convergence. In the negative-gradient prototype (e.g., Figure 8B, left), the overall average phase was negative and the average phase of each phase curve was negatively correlated with the size of frequency convergence (more negative for larger frequency convergence). In the positive-gradient prototype (e.g., Figure 8B, middle and right), the overall average phase was positive and the average phase of each phase curve was positively correlated with the size of frequency convergence (more positive for larger frequency convergence). Thus, in the two-dimensional space defined by the overall average phase on the x-axis and the average-phase-vs-convergence-size correlation on the y-axis with the lines x=0 and y=0 dividing the space into four quadrants, the negative-gradient prototype is represented in the lower-left quadrant and the positive-gradient prototype is represented in the upper-right quadrant. We excluded from this analysis the phase curves for the largest frequency convergence (i.e., convergence of all sites), which tended to be noisy due to the relatively rare occurrences of the maximal convergence (Figure 4).

As shown in Figure 8A, 8C, and 8E (minimal visual stimulation) and Figure 9A and 9C (natural viewing), most participants indeed populated either the lower-left quadrant (red-shaded rectangles) or the upper-right quadrant (blue-shaded rectangles). The average phase-gradient patterns are shown in Figure 8B, 8D, and 8F (minimal visual stimulation) and Figure 9B and 9D (natural viewing) for each quadrant that were populated by five or more participants. To statistically confirm that most participants populated the lower-left and upper-right quadrants in all cases, we conducted *χ*^2^(1) tests comparing the number of participants populating these quadrants versus the number of participants populating the remaining two quadrants.

**Figure 9.**
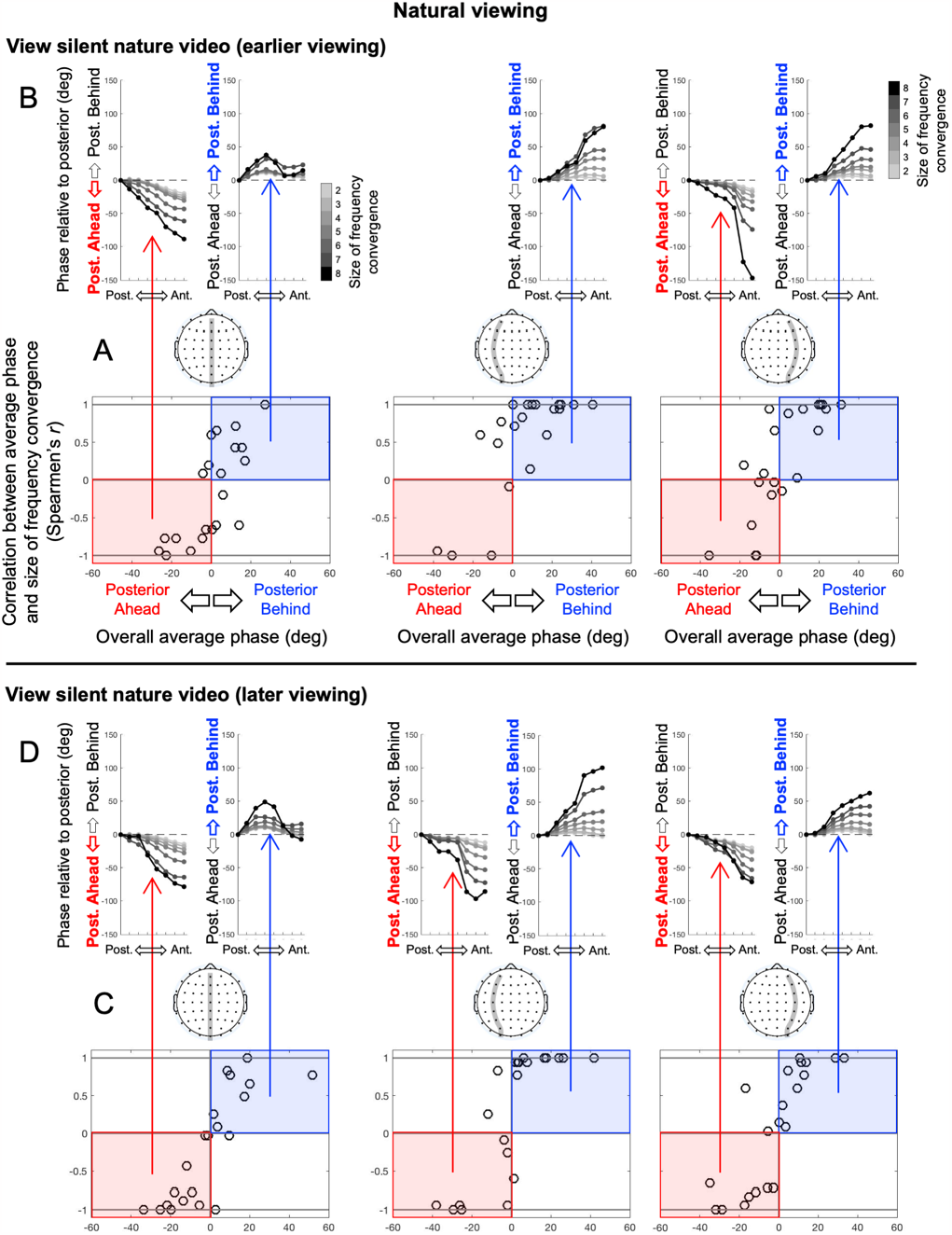
Two general types of posterior-to-anterior phase gradients observed during inter-site oscillation-frequency convergences in natural viewing: the earlier (A and B) and later (C and D) viewing of a silent nature video. The formatting of this figure is the same as in Figure 8. Again, most participants populated either the lower-left quadrant (red rectangles in A and C) characterized by generally negative posterior-to-anterior phase gradients that became steeper (more negative) as the size of frequency convergence increased (red arrows in B and D), or the top-right quadrant (blue rectangles in A and C) characterized by generally positive posterior-to-anterior phase gradients that became steeper (more positive) as the size of frequency convergence increased (blue arrows in B and D). However, unlike under minimal visual stimulation (Figure 8), the negative and positive gradients were not consistently spatially organized as participants were relatively evenly distributed between the lower-left and upper-right quadrants regardless of the three anterior-posterior axes.

The results were significant in all 15 cases (five behavioral conditions times three anterior-posterior axes). For the rest-with-eyes-closed condition, *χ*^2^(1) = 16.667, *p* < 0.0001, and Cramer’s *V* = 0.833 for the midline axis, *χ*^2^(1) = 13.500, *p* < 0.0003, and *V* = 0.750 for the left-hemisphere axis, and *χ*^2^(1) = 10.667, *p* < 0.002, and *V* = 0.667 for the right-hemisphere axis. For its replication, *χ*^2^(1) = 16.667, *p* < 0.0001, *V* = 0.833 for the midline axis, *χ*^2^(1) = 24.000, *p* < 0.0001, and *V* = 1.000 for the left-hemisphere axis, and *χ*^2^(1) = 24.000, *p* < 0.0001, and *V* = 1.000 for the right-hemisphere axis. For the rest-with-eyes-open-in-the-dark condition, *χ*^2^(1) = 8.167, *p* < 0.005, and *V* = 0.583 for the midline axis, *χ*^2^(1) = 16.667, *p* < 0.0001, and *V* = 0.833 for the left-hemisphere axis, and *χ*^2^(1) = 24.000, *p* < 0.0001, and *V* = 1.000 for the right-hemisphere axis. For the silent-nature-video condition (earlier viewing), *χ*^2^(1) = 3.857, *p* < 0.05, and *V* = 0.429 for the midline axis, *χ*^2^(1) = 10.714, *p* < 0.002, and *V* = 0.714 for the left-hemisphere axis, and *χ*^2^(1) = 8.48, *p* < 0.005, and *V* = 0.619 for the right-hemisphere axis. For the silent-nature-video condition (later viewing), *χ*^2^(1) = 13.762, *p* < 0.0003, and *V* = 0.810 for the midline axis, *χ*^2^(1) = 10.714, *p* < 0.002, and *V* = 0.714 for the left-hemisphere axis, and *χ*^2^(1) = 13.762, *p* < 0.0003, and *V* = 0.810 for the right-hemisphere axis. Thus, the negative and positive gradients resembling the prototypes were prevalent across all three anterior-posterior axes in all behavioral conditions.

Interestingly, under minimal visual stimulation (especially with the eyes closed) the negative- and positive-gradient patterns were spatially organized. The negative-gradient patterns consistently formed along the midline axis (most participants populating the lower-left quadrant in the left panels in Figure 8A and 8C, and to some degree in 8E), whereas the positive-gradient patterns formed along the left- and right-hemisphere axes (most participants populating the upper-right quadrant in the middle and right panels in Figure 8A, 8C, and 8E). To confirm this observation, we conducted *χ*^2^(1) tests evaluating the hypothesis that the lower-left quadrant (relative to the upper-right quadrant) was more populated for the midline axis whereas the upper-right quadrant (relative to the lower-left quadrant) was more populated for the left- and right-hemisphere axes (combined). The results were significant for all three conditions under minimal visual stimulation: *χ*^2^(1) = 43.608, *p* < 0.0001, and Cramer’s *V* = 0.825 for the rest-with-eyes-closed condition, *χ*^2^(1) = 29.703, *p* < 0.0001, and *V* = 0.651 for its replication, and *χ*^2^(1) = 16.010, *p* < 0.0001, and *V* = 0.496 for the rest-with-eyes-open-in-the-dark condition. This confirms that the negative-gradient patterns were prevalent along the midline axis whereas the positive-gradient patterns were prevalent along the left- and right-hemisphere axes under minimal visual stimulation.

We further confirmed that the phase gradients were positively correlated between the left- and right-hemisphere axes whereas they were negatively correlated between the left- and right-hemisphere axes and the midline axis. To obtain a single gradient profile per anterior-posterior axis per participant, we averaged the gradients across frequency-convergence sizes. For the rest-with-eyes-closed condition and its replication, the left- and right-hemisphere gradients were significantly positively correlated (Fisher-*Z* transformed Pearson’s *r, M* = 1.544, *SD* = 1.007, *t*(23) = 7.511, *p* < 0.0001, and *d* = 1.533 for the rest-with-eyes-closed condition; *M* = 1.131, *SD* = 0.871, *t*(23) = 6.366, *p* < 0.0001, and *d* = 1.299 for its replication), and the left- and right-hemisphere gradients (averaged) were significantly negatively correlated with the midline gradient (*M* = –1.016, *SD* = 1.273, *t*(23) = –3.910, *p* < 0.0008, and *d* = –0.798 for the rest-with-eyes-closed condition; *M* = –0.855, *SD* = 1.177, *t*(23) = –3.558, *p* < 0.002, and *d* = –0.726 for its replication). For the rest-with-eyes-open-in-the-dark condition, the positive correlation between the left- and right-hemisphere gradients was significant (*M* = 1.024, *SD* = 1.033, *t*(23) = 4.857, *p* < 0.0001, and *d* = 0.991), but the negative correlation between the left- and right-hemisphere gradients (averaged) and the midline gradient did not reach significance (*M* = –0.376, *SD* = 1.290, *t*(23) = –1.428, *p* = 0.167, and *d* = –0.292). Thus, the spatial organization of the negative and positive gradients was robust while participants rested with their eyes closed, but it somewhat weakened when participants opened their eyes even under minimal visual stimulation.

The spatial organization dissipated in natural viewing. Participants relatively evenly populated the lower-left quadrant (indicative of the negative-gradient patterns) and the upper-right quadrant (indicative of the positive-gradient patterns) regardless of the axes, the midline axis (first panels in Figure 9A), the left-hemisphere axis (second panels in Figure 9A), or the right-hemisphere axis (third panels in Figure 9A). The *χ*^2^(1) tests used above to confirm the spatial organization were non-significant: *χ*^2^(1) = 0.918, *p* = 0.338, and Cramer’s *V* = 0.137 for the silent-nature-video (earlier viewing) condition, and *χ*^2^(1) = 1.520, *p* = 0.218, and *V* = 0.165 for the silent-nature-video (later viewing) condition.

While the opposing gradients were spatially organized only under minimal visual stimulation, for most participants, the maximum inter-axis correlation of the phase gradients (averaged across frequency-convergence sizes) was positive, indicating that two of the three axes had the same gradient direction, while the minimum correlation was negative, indicating that one axis had the opposite gradient direction. This patten was obtained from 20/24 of participants or 83% (*χ*^2^(1) = 10.667, *p* < 0.002, and Cramer’s *V* = 0.667) for the rest-with-eyes-closed condition, 20/24 or 83% (*χ*^2^(1) = 10.667, *p* < 0.002, and *V* = 0.667) for its replication, 19/24 or 79% (*χ*^2^(1) = 8.167, *p* < 0.005, and *V* = 0.583) for the rest-with-eyes-open-in-the-dark condition, 16/21, or 76% (*χ*^2^(1) = 5.762, *p* < 0.02, and *V* = 0.524) for the silent-nature-video (earlier viewing) condition, and 15/21 or 71% (*χ*^2^(1) = 3.857, *p* < 0.05, and *V* = 0.429) for the silent-nature-video (later viewing) condition. These ratios did not differ significantly across the conditions, *χ*^2^(4) = 0.163, *p* = 0.997, and *V* = 0.028. Thus, for most participants, the opposing gradient patterns were present across the three axes in all conditions, except that they were idiosyncratically distributed across the three axes in natural viewing, but were consistently spatially organized under minimal visual stimulation (especially when the eyes were closed).

These phase-gradient patterns remained relatively stable over time, yielding phase-clustering values (PCV’s, also known as phase-locking values, PLV’s: the modulus of time-averaged complex phase differences ranging between 0 and 1) of ∽0.7 above chance levels (computed with the assumption of random sampling from a uniform angular distribution, well approximated by a power function of the number of instances). The stability of the phase gradients (PCV’s of sequential pairwise phase differences) was relatively constant along the midline axis (left column in Figure 10A). Along the left- and right-hemisphere axes, the stability varied with some consistency (left column in Figure 10B and 10C), and the variations were similar for different sizes of frequency convergence (right column in Figure 10B and 10C). Future research may interpret these consistent variations. Importantly, the overall stability of the phase gradients was not substantially affected by the size of frequency convergence (right column in Figure 10A, 10B, and 10C), except that the stability was somewhat reduced for larger frequency convergences along the left- and right-hemisphere axes in natural viewing (bottom two rows in the right column in Figure 10B and 10C). Overall, oscillation-frequency convergences did not have substantial effects on the stability of phase relations.

**Figure 10.**
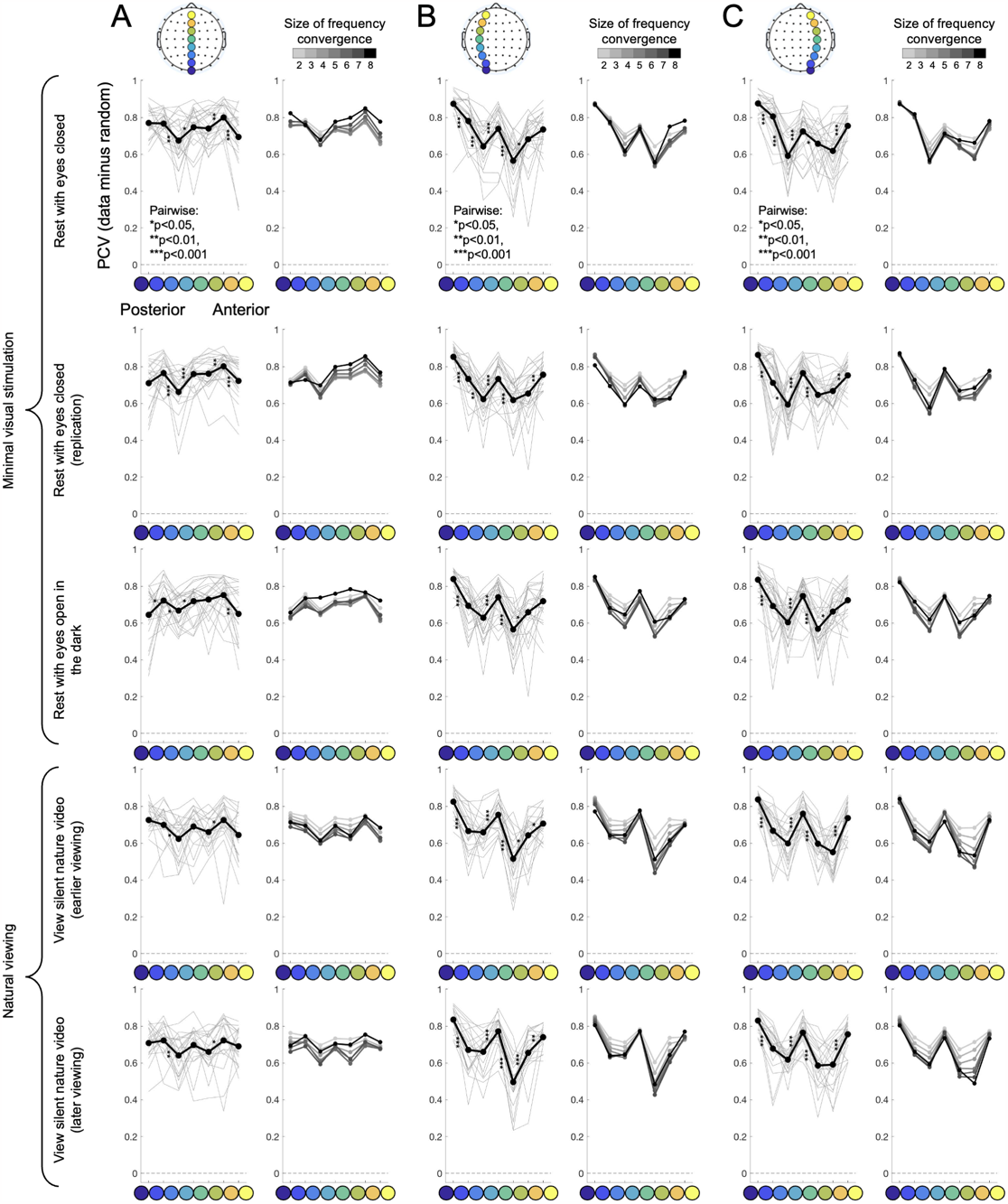
Phase-clustering values (PCV’s, also known as phase-locking values or PLV’s) computed for sequential pairs of anterior-posterior sites (cooler to warmer colors) along the midline (**A**), left-hemisphere (**B**), and right-hemisphere (**C**) axes for the five behavioral conditions (rows). **Left panels**. Average PCV’s (black curves) and those for individual participants (gray curves). Asterisks indicate statistical significance for comparisons between successive pairs of values. **Right panels**. Average PCV’s shown separately for frequency-convergences of two through eight sites (coded with lighter to darker gray). PCV’s for random angular distributions of the corresponding numbers of instances have been subtracted so that the value of zero (dashed lines) indicates chance levels.

While we conducted our oscillation-frequency convergence analyses in the extended alpha range (5-15 Hz), the question remained whether large frequency convergences occurred evenly in this range or in a specific subrange. We thus computed average frequency-convergence size as a function of frequency by taking each participant’s oscillation-frequency-(row)-by-time-(column) flow-convergence matrix (per anterior-posterior axis per condition) and averaging the convergence sizes (omitting zeros where no oscillations occurred at any sites) across time for each oscillation frequency. To control for any influences of time independent spectral characteristics at individual sites (see above), we subtracted the values obtained from the corresponding time-shuffled controls (averaged across 20 versions). To avoid the analysis being unduly influenced by participants who frequently yielded large convergence sizes, we normalized each participant’s average-convergence-size-by-frequency curve by dividing each value by the average value across all frequencies (per anterior-posterior axis per condition). As shown in Figure 11, the size of oscillation-frequency convergence was substantially elevated in the conventional alpha range (8-12 Hz) for all anterior-posterior axes in all conditions.

**Figure 11.**
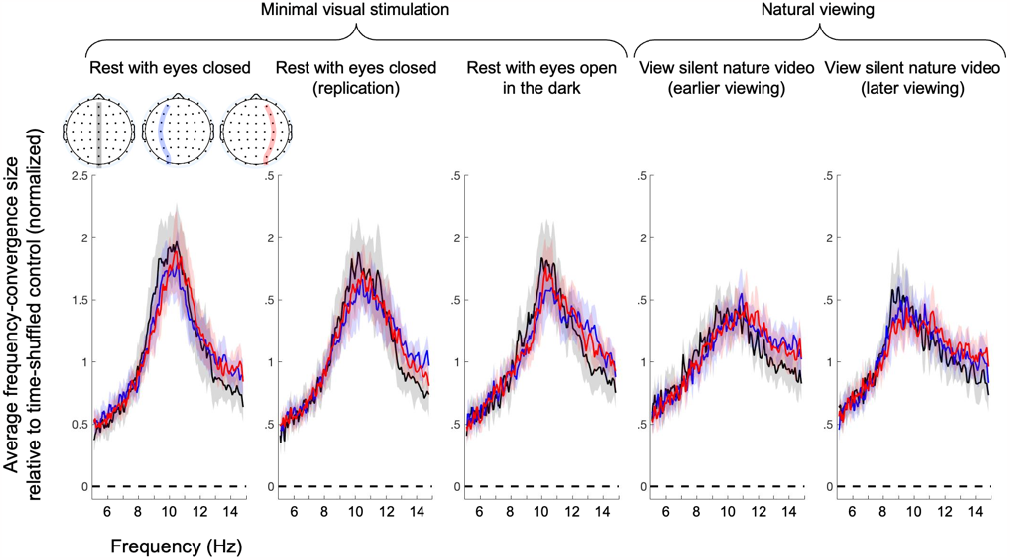
Time-averaged size of anterior-posterior frequency convergence as a function of oscillation frequency (Hz) along the midline (black), the left hemisphere (blue) and right hemisphere (red) axes for the five behavioral conditions. The values are relative to those obtained with the corresponding time-shuffled controls (averaged across 20 versions). The curve from each participant was normalized by dividing each value by the average value across all frequencies. Note that oscillation-frequency convergences were substantially elevated in the conventional alpha range (8-12 Hz) in all cases. The shaded areas represent the 95% confidence intervals (computed with participants as the random effect).

Thus, the results reported here primarily reflect oscillation-frequency convergences occurring in the alpha band.

## Discussion

Prior work on oscillatory neural interactions typically examined inter-regional phase coherence in specific frequency bands (without identifying oscillation frequencies) to study frequency-specific functional networks. We took a complementary approach of tracking regional oscillation frequencies to investigate how brain regions dynamically converge their oscillation frequencies to stably interact with one another via phase locking. Given the known functional relevance of anterior-posterior neural interactions (e.g., de Pasquale et al., 2012; Gonzalez-Castillo and Bandettini, 2018; Lobier et al., 2018; Marzetti et al., 2019; Mashour et al., 2020; Reinhart and Nguyen, 2019), the presence of various neuroanatomical and neurophysiological feature gradients along the anterior-posterior axis (e.g., Burt et al., 2018; Huntenburg et al., 2018; Mahjoory et al., 2020), and the relevance of theta, alpha, and low-beta bands in organizing gamma-oscillation activity (e.g., Bahramisharif et al., 2013; Esghaei et al., 2022), controlling information flows (e.g., Jensen et al., 2014; Muller et al., 2018; Zhang et al., 2018), and mediating perceptual, attentional, and memory processes (e.g., Bonnefond and Jensen, 2015; Busch and VanRullen, 2010; Fries, 2015; Harris et al., 2018; Klimesch, 2012; Lobier et al., 2018; Mathewson et al., 2010; Palva and Palva, 2007; Reinhart and Nguyen, 2019), we examined the dynamics of oscillation-frequency convergences along anterior-posterior axes in an extended alpha range (5-15 Hz) while participants rested with their eyes open or closed. Oscillation-frequency convergences were examined along three anterior-posterior axes (each containing eight sites), along the midline and within each hemisphere.

Oscillation-frequency convergences primarily occurred in the conventional alpha band (8-12 Hz). The instances of large frequency convergences (involving four or more sites out of eight) were elevated for all participants in all behavioral conditions relative to the time-shuffled controls, suggesting that oscillation frequencies dynamically converge along anterior-posterior axes over and above any tendencies for dominant oscillations in relatively narrow frequency bands to randomly coincide. The probability of an additional site joining a frequency convergence increased as more sites frequency-converged, suggesting that synergistic interactions drive oscillation-frequency convergences. The oscillatory power at participating sites increased as more sites frequency-converged (especially as five of more sites converged), suggesting that the synergistic interactions are mediated by regional synchronizations, being boosted by inter-regional frequency matching, facilitating the entrainment of additional regions. This interpretation is consistent with previous reports of elevated oscillatory power in brain regions engaged in strong and/or extensive phase locking with other regions (e.g., Moon et al., 2015; Tewarie et al., 2019).

Given our results suggest that synergistic interactions drive spontaneous anterior-posterior oscillation-frequency convergences, what functions might they serve? Although we did not include behavioral experiments to compare frequency convergences with behavioral task performances, some inferences can be made from the two patterns of anterior-posterior phase gradients that we observed during oscillation-frequency convergences. One was a negative gradient from posterior to central sites (generally plateauing from central to anterior sites)—the posterior-ahead gradient. The other was a positive gradient from posterior to central sites (generally plateauing from central to anterior sites)—the posterior-behind gradient. These gradients remained relatively stable over a 5-min period, yielding the average PCV’s of ∽0.7 above chance levels.

The posterior-ahead gradient is consistent with the interpretation (though it does not necessarily imply) that information flows out of posterior cortex in the sense that oscillations ahead in phase may be causing those behind in phase. Similarly, the posterior-behind gradient is consistent with the interpretation that information flows to posterior cortex. These gradients generally became steeper as the size of frequency convergence increased. A steeper phase gradient would suggest stronger directionality of information flow provided that the stability of the phase gradient was not reduced by the increased slope. The stability (in PCV’s relative to chance levels) was not systematically affected by the size of frequency convergence along the midline, though it somewhat reduced with larger frequency convergences in the left and right hemispheres, especially in natural viewing. Thus, the steeper slopes associated with larger frequency convergences suggest stronger directionality of the posterior-ahead and posterior-behind phase gradients at least along the midline as well as in both hemispheres under minimal visual stimulation. In other words, frequency convergences enhance the directionality of information flows to and from posterior regions especially along the midline, but potentially also in both hemispheres especially under minimal visual stimulation.

The anterior-posterior phase gradients we observed were based on phase relations that remained stable over relatively long (∽5 min) periods. Further, those phase gradients were piece-wise consistent whether only a few sites or most sites frequency-converged except that the gradients became steeper when more sites frequency-converged. As such, the posterior-behind and posterior-ahead gradients observed here do not imply macroscopic traveling waves, which have been demonstrated to generally propagate from posterior to anterior when the eyes are open and from anterior to posterior when the eyes are closed (e.g., Alamia and VanRullen, 2019; Alexander et al., 2013; Zhang et al., 2018). Further, one of the primary mechanisms of traveling waves is a linear spatial gradient of oscillation frequencies, wherein a wave travels from higher-frequency regions to lower-frequency regions (e.g., Ermentrout and Kleinfeld, 2001). In contrast, the stable anterior-posterior phase gradients we observed occurred when participating regions oscillated at matched frequencies. Thus, macroscopic traveling waves of various amplitudes likely propagate in either direction in different moments against the background of the stable phase gradients we observed that are enhanced by oscillation-frequency convergences.

How did anterior-posterior oscillation-frequency convergences depend on the midline versus within-hemisphere axes and behavioral conditions? The prevalence of oscillation-frequency convergences, the increased probability of growing in convergence with increased size of frequency convergence, and the increased oscillatory power with the increased size of frequency convergence, were relatively independent of the location of the anterior-posterior axis and behavioral conditions (except that they all somewhat diminished in natural viewing relative to under minimal visual stimulation). This suggests that the synergistic interactions that likely drive spontaneous oscillation-frequency convergences are ubiquitous. In contrast, the manner of spreading of frequency convergences and the occurrences of the posterior-behind and posterior-ahead phase gradients depended on the location of the anterior-posterior axis and behavioral conditions.

Frequency convergences spread differently along the midline versus within each hemisphere. Along the midline, the anterior-most and posterior-most sites joined oscillation-frequency convergences with lower likelihood, with the pattern consistent across all behavioral conditions. This suggests that anterior-posterior frequency convergences along the midline tend to spread from any site except the anterior-most and posterior-most sites. In the left and right hemispheres, more anterior sites joined oscillation-frequency convergences with progressively lower likelihoods. This trend was stronger under minimal visual stimulation than in natural viewing. This suggests that anterior-posterior frequency convergences within each hemisphere tend to spread from posterior to anterior regions especially under minimal visual stimulation.

The posterior-ahead and posterior-behind phase gradients were consistently organized under minimal visual stimulation. When participants rested with their eyes closed, the posterior-ahead gradients predominantly formed along the midline axis whereas the posterior-behind gradients predominantly formed along the left- and right-hemisphere axes. This spatial organization weakened when participants opened their eyes even in the dark. The opposing gradients distributed idiosyncratically across the three anterior-posterior axes in natural viewing. This suggests that most people, when they close their eyes, default to the pattern of information flow where information carried in the alpha band (8-12 Hz) flows to posterior regions within each hemisphere and flows out of posterior regions along the midline. Future research may investigate how closing eyes induces these spatially structured anterior-posterior information flows and how they influence information processing.

In summary, our results suggest that synergistic interactions drive inter-regional oscillation-frequency convergences in the alpha band, wherein regional synchronizations, being boosted by inter-regional frequency matching, facilitate the entrainment of additional regions. Our results also suggest that inter-regional oscillation-frequency convergences generate stable posterior-behind gradients (potentially facilitating information flows to posterior regions) and posterior-ahead phase gradients (potentially facilitating information flows out of posterior regions), which increase in directionality as more sites frequency-converge and spatially organize when the eyes are closed. Future research may investigate the behavioral relevance of these characteristics of anterior-posterior oscillation-frequency convergences by relating them to task performances.

